# The landscape of somatic mutation in sporadic Chinese colorectal cancer

**DOI:** 10.1101/155671

**Authors:** Zhe Liu, Chao Yang, Xiangchun Li, Wen Luo, Bhaskar Roy, Teng Xiong, Xiuqing Zhang, Huanming Yang, Jian Wang, Zhenhao Ye, Yang Chen, Jinghe Song, Shuai Ma, Yong Zhou, Min Yang, Xiaodong Fang, Jie Du

## Abstract

Colorectal cancer is the fifth prevalent cancer in China. Nevertheless, a large-scale characterization of Chinese colorectal cancer mutation spectrum has not been carried out. In this study, we have performed whole exome-sequencing analysis of 98 patients’ tumor samples with matched pairs of normal colon tissues using Illumina and Complete Genomics high-throughput sequencing platforms. Canonical CRC somatic gene mutations with high prevalence (>10%) have been verified, including *TP53, APC, KRAS, SMAD4, FBXW7* and *PIK3CA. PEG3* is identified as a novel frequently mutated gene (10.6%). *APC* and Wnt signaling exhibit significantly lower mutation frequencies than those in TCGA data. Analysis with clinical characteristics indicates that *APC* gene and Wnt signaling display lower mutation rate in lymph node positive cancer than negative ones, which are not observed in TCGA data. *APC* gene and Wnt signaling are considered as the key molecule and pathway for colorectal cancer initiation, and these findings greatly undermine their importance in tumor progression for Chinese patients. Taken together, the application of next-generation sequencing has led to the determination of novel somatic mutations and alternative disease mechanisms in colorectal cancer progression, which may be useful for understanding disease mechanism and personalizing treatment for Chinese patients.

## 1 INTRODUCTION

Colorectal cancer (CRC) is the third most prevalent type of cancer worldwide [1, 2]. CRC has nearly 4 millions new cases and caused 2 millions death in China each year [1, 2]. In the last decade, its incidence has increased constantly in China due to the diet and living habit change [1]. CRC patients usually attain promising clinical outcomes of early diagnosis, however, most of them failed to detect the tumor until a late stage. Around 60% of CRC reside on rectum in Chinese patients, while 40% occurs on rectum in Caucasian patients [3].

Currently, various genomic approaches have been devoted to investigate the molecular mechanisms of CRC initiation and progression. A pioneering study identified several somatic mutations in colorectal cancer, such as *TP53, KRAS, APC, PIK3CA* and *FBXW7* [4]. A later TCGA study has shed light on utilizing genomic data to elucidate mutation landscape of human CRC, and several novel somatic mutations were identified as well [5]. A follow-up study revealed several novel somatic mutations, such as *TCF7L2, TET2, TET3* and *ERBB3*, and illustrated possible treatment plan for colorectal cancer [6]. Whole exome sequencing (WES) study on American-African patients discovered significant different somatic mutation genes, indicating alternative disease mechanisms in patients with different ethnic background [7]. Moreover, investigations on Iranian and Japanese patients uncover different somatic gene mutations and alternative mutation frequencies than Caucasian counterparts [8, 9]. Therefore, a whole exome sequencing of Chinese patients is essential for novel somatic gene mutation spectrum characterization, which may consequently change our understanding of disease etiology and precision medicine management.

The most relevant study of Chinese CRC using exome sequencing has used whole exome sequencing for the first CRC cancer prognostic study [10]. A few novel somatic mutation genes, such as *CDH10, FAT4* and *DOCK2*, are reported to be susceptible driver mutations. Notwithstanding, due to its limited sample size (22 samples) and low sequencing coverage (< 60X) strategy, its power for novel Chinese CRC gene discovery and mutation frequency characterization is limited for this high immune and stromal cell infiltration tumor [11]. Up to now, a genome-wide somatic gene mutation and frequency data are largely unknown for Chinese patients. Also, little was discussed towards analysis of the association between gene mutations and clinical characteristics for Chinese patients.

In this study, we used whole exome sequencing technology to study sporadic Chinese CRC, a single type of CRC taking for around 65-85% of the total CRC patients. WES of 98 sporadic Chinese CRC patients’ samples with matched controls is sequenced by Illumina and Complete Genomics platforms. First, we compared the somatic mutations and mutation frequency with TCGA data. Second, we carried out pathway analysis and compared them with TCGA data. Finally, we analyzed clinical characteristics by their association with somatic mutated genes.

## 2 RESULTS

### 2.1 Sequencing statistics

To identify genetic variants involved in CRC, we performed whole-exome sequencing of the tumor and matched controls on genomic DNA from 40 (Second affiliated Hospital of Nanchang University) and 58 patients (Wenzhou Medical University) by Illumina HiSeq2000 (ILMN) and Complete Genomics (CG) Blackbird sequencers, respectively. The Illumina and CG platforms attained 117X and 168X average coverage for exon captured regions as shown in Figure 1a. All of the samples achieved 10X and 20X coverage rate more than 95% and 90% of the genome for Illumina and CG platforms. A description of sequencing and mutation statistics can be found in Supplementary Table S2.

**Fig. 1.**
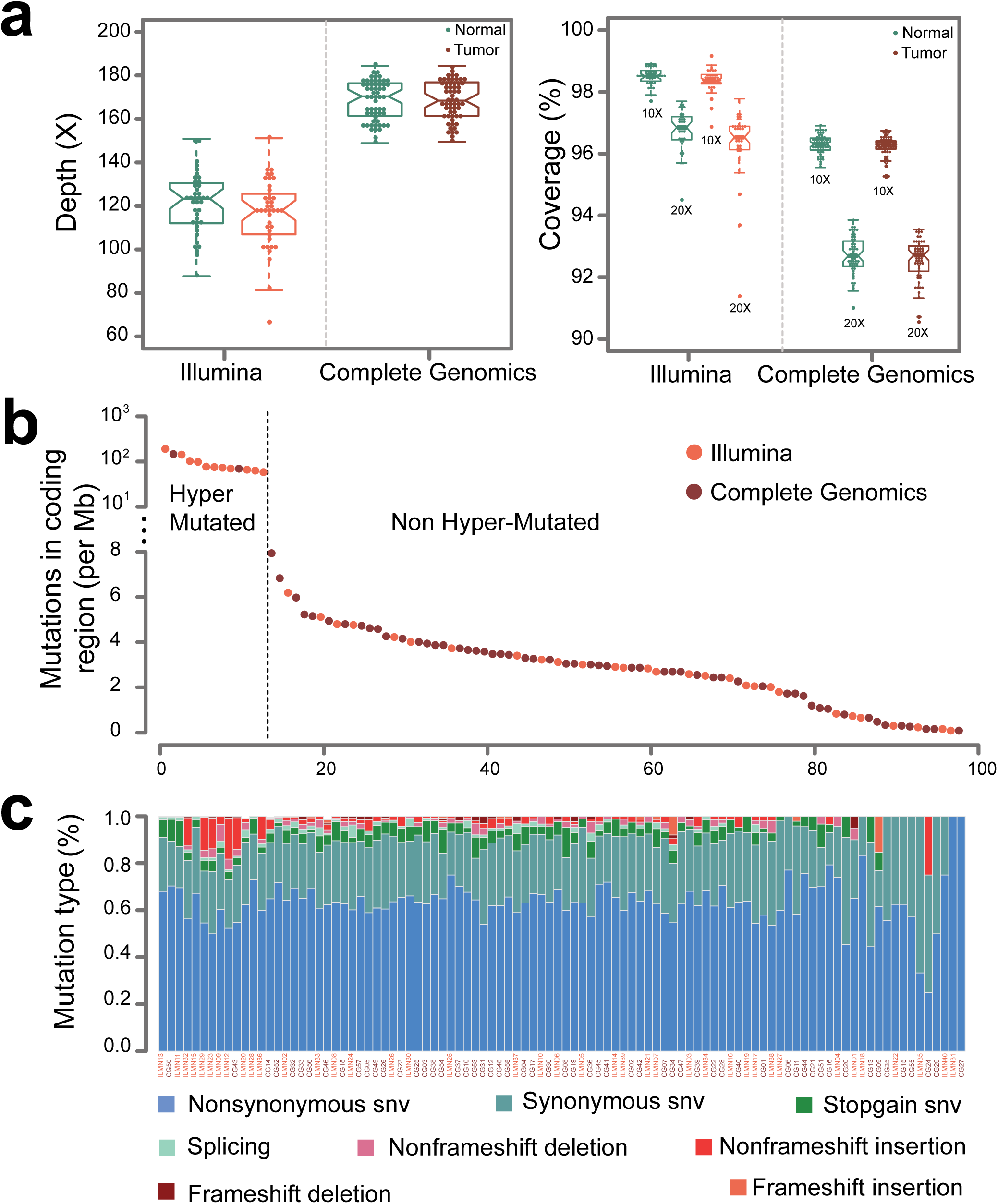
Sequencing statistics for Illumina and CG platforms. (a) This graph displays the average sequencing depth and exome coverage percentage with >10X and >20X for Illumina and Complete Genomics platforms. The left chart describes the average number of reads aligned to human reference genome hg19. The average depth can be computed from the length of the original genome (G), the number of reads (N), and the average read length (L) as {N*L/G}. (b) This chart illustrates the mutations in coding regions for Illumina and Complete Genomics platforms. The dash line is used to separate samples into hyper-mutated and regularly mutated ones. (c) A display of the various categories of mutations across samples is shown for SNVs (non-synonymous SNV, synonymous SNV, stopgain SNV and splicing) and InDels (non-frameshift deletion, non-frameshift insertion, frameshift deletion and frameshift insertion).

It was reported that CG platform has high concordance rate for SNV detection with Illumina platform [12]. However, it is necessary to evaluate system bias from two sequencers by their ability to discover somatic mutations. This was verified by no segregation of mutation numbers in coding region from two platforms (Figure 1b). Also, no obvious differences were observed between the mutation counts (p=0.29, Wilcoxon rank test). Moreover, mutation count of each type of SNV (non-synonymous SNV, synonymous SNV, stop gain SNV, splicing SNV) and InDel (non-frame shift deletion InDel, non-frame shift insertion InDel, frame shift deletion InDel, frame shift insertion InDel) does not show statistical differences between the two platforms. Taken together, it is reasonable to merge these two datasets for further study. Mutation rates of tumors are around 3/Mb for each sample (Figure 1c), which is consistent with previous western and Chinese CRC whole exome sequencing results [5, 10, 13].

Overall, 13 patients have considerably higher mutation rate (Figure 1b), and we treat them as hypermutated samples. CG and Illumina platforms are able to identify 2.85M and 2.95M mutations in coding region for the non-hypermutated samples. A report of the detected SNVs and InDels are shown in Supplementary Table S3 and S4.

### 2.2 Somatic mutational spectrum

In general, dominant somatic SNVs in colorectal cancer are *CpG->T mutations as shown in Figure 2a, consistent with that of TGCA (Figure 2b). To investigate the origin of somatic mutations, we clustered samples into subgroups based on the six mutation types (Supplementary Appendix A). We stratified the patients into hypermutated and regularly mutated subgroups for Chinese data. Similar results are also derived from TCGA data, indicating the consistency of the clustering results. The differences between the mutation spectra of the two subgroups are quite obvious. Mutations in regularly mutated samples are mainly *CpG->T, while hypermutated samples are dominated by three mutation peaks at TCT>TAT, TTT>TGT and TCG>TTG.

**Fig. 2.**
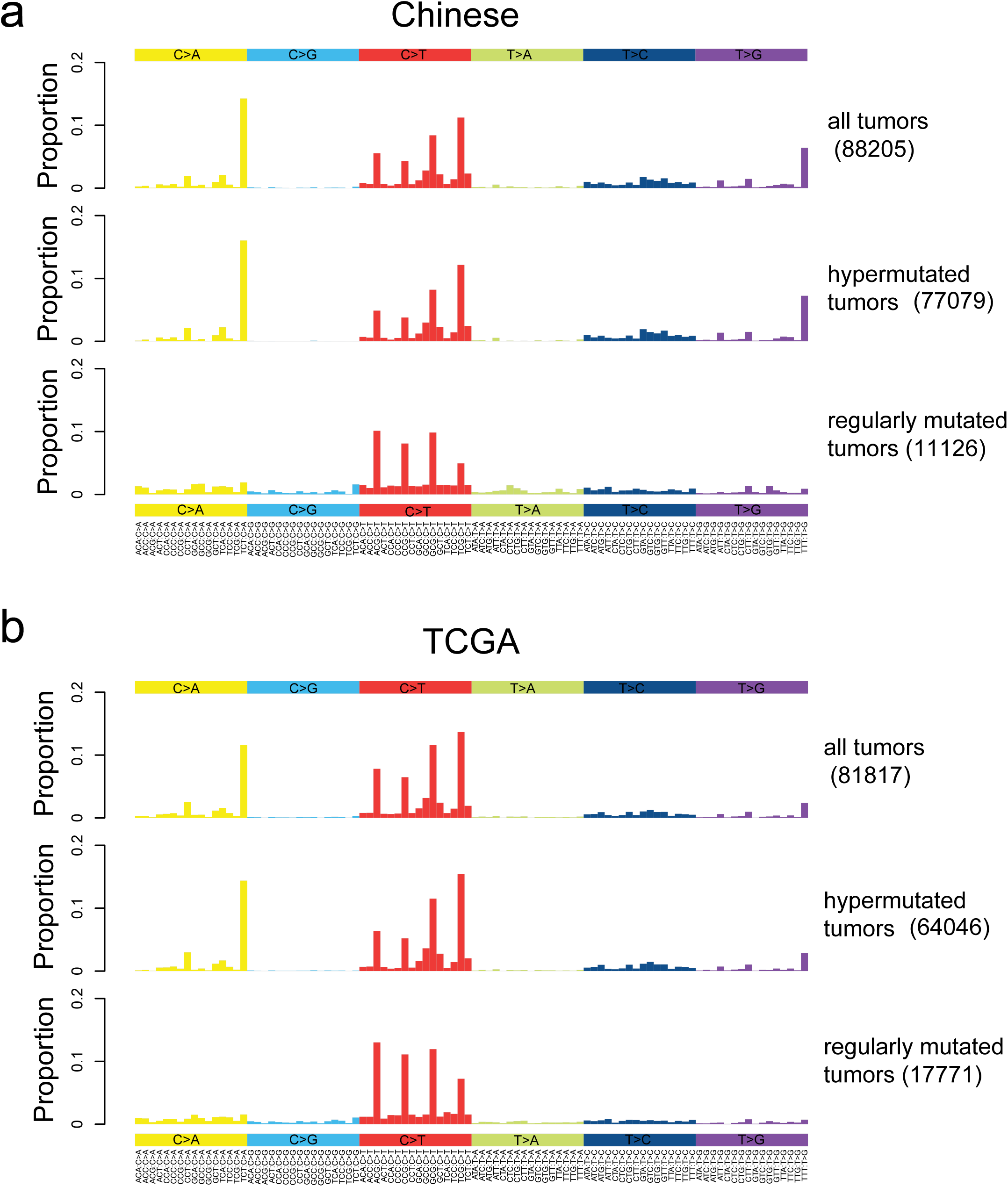
Somatic mutation spectrums for all, hypermutated and regularly mutated samples. Base substitutions are divided into six categories, and each is represented by a color. The 16 possible flanking nucleotide types for each category are then plotted on the horizontal axis. The vertical axis shows the proportion of somatic mutations of each type. (a). Mutation spectrums for all, hypermutated and regularly mutated Chinese samples. (b). Mutation spectrums for all, hypermutated and regularly mutated TCGA samples.

### 2.3 Somatic mutation

We performed somatic gene mutation analysis of the hypermutated samples due to its unique tumorigenisis process and clinical outcome. It is known in the previous research that most of the microsatellite instability (MSI) tumors are hypermutated [5]. This is also validated in our study, and 8 out of 11 the hypermuated samples are MSI. The rest 3 hyper-mutated and microsatellite stability (MSS) samples displays mutations on *MSH4* and *POLE*. Mutations on *MSH4* and *POLE* may interrupt DNA binding and DNA replication, both of which will induce ultra-high mutation numbers. Moreover, canonical CRC genes, including *APC, PIK3CA, MSH6* and *FAT4* are observed in mutation status. It is interesting to discover *TP53* in lower mutation prevalence (2 out of 13) for hypermutated samples, which may indicate alternative disease mechanisms for hypermutated samples. A report of the prevalently mutated genes is shown in Supplementary Table S5.

We then discuss the somatic mutations in non-hypermutated samples. In general, identification of driver mutations from passenger mutations is a challenging task due to the following reasons: a) CRC is a type of cancer with high immune and stromal cells infiltration and somatic mutation signal will be diminished [14]. b) Patients may have sub-clone with different somatic mutated genes, and a mixture of the two or more sub-clone will further decrease somatic mutation signal [15, 16]. c) Disease etiology for a fraction of patients may be induced by different somatic mutations and a long tail of genes was postulated to explain the CRC initiation and progression of entire population [11]. d) Consistency between different significantly mutated genes algorithms are not perfect, and genes shown up as statistical significant results from one algorithm may disappear in another one [17, 18].

In this study, we used the following rules to discuss the susceptible driver somatic mutations. 1) We used MutsigCV, one of the most widely used algorithms, for significantly mutated gene (SMG) detection [19]. 2) We reported prevalently mutated somatic genes (>5% of the entire population) due to its possibility as diagnostic and treatment targets on population level. 3) We discussed the mutated genes supported by additional literatures. 4) We used two gene expression data (GSE50760 and GSE18105) from GEO database as gene expression evidences. Genes with log fold change>1 and adjusted p-value<0.05 between tumors and matched controls are considered as differentially expressed genes [20, 21]. A report of these genes was presented in Supplementary Table S6 and Figure 3a. It should be mentioned that the two platforms achieve high consistency in terms of significant somatic mutated gene identification and gene mutation frequency characterization.

**Fig. 3.**
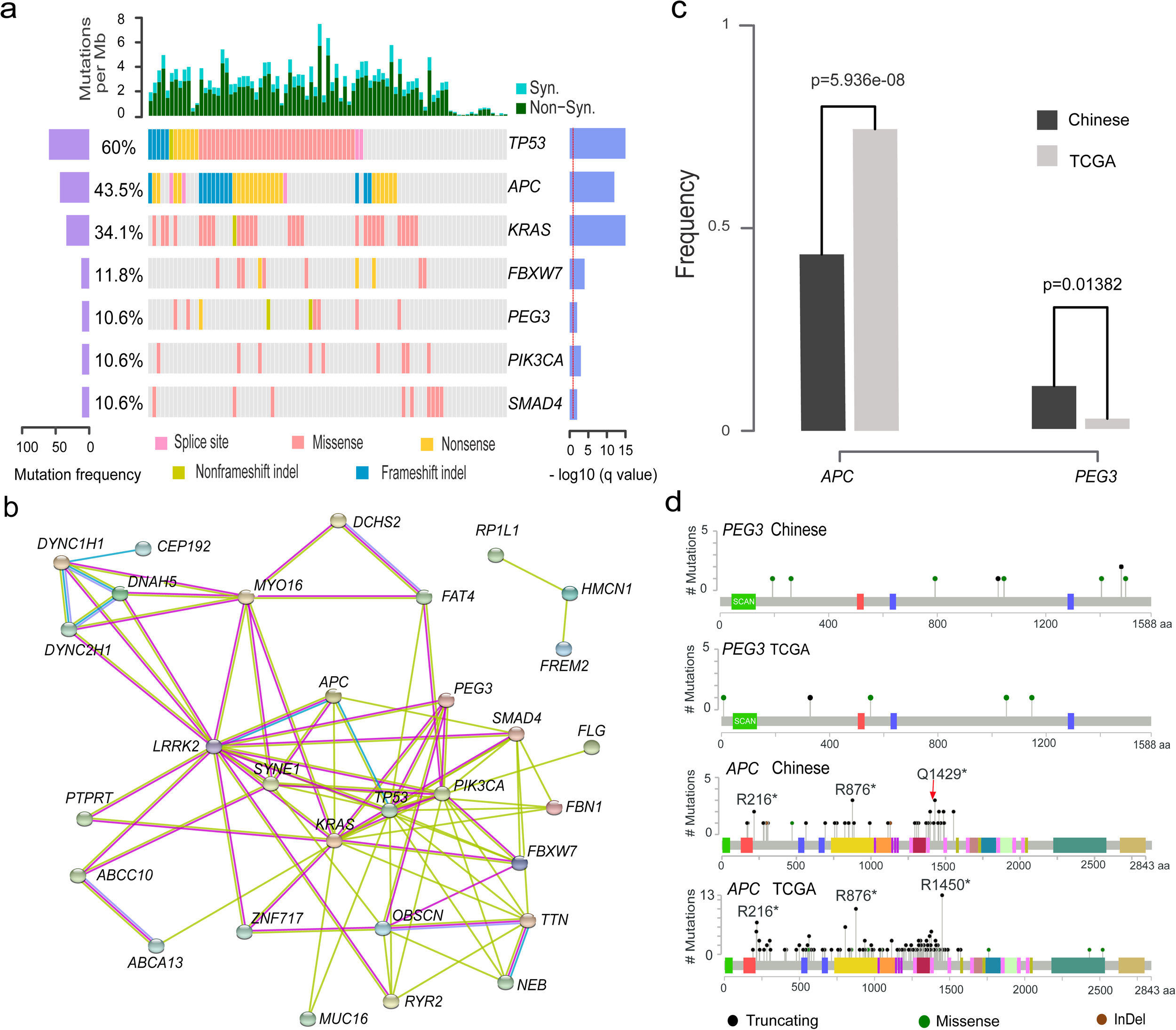
Illustration of prevalently somatic mutated genes. (a). Illustration of significantly mutated genes in non-hypermutated samples. The left axis shows mutation frequencies in 85 non-hypermutated samples. The right axis indicates the –log10 transformed q-value score from MutSigCV. (b) Protein interaction network from String database of mutated genes with >5% frequency. Green, red and blue edges represent the evidence from text-mining, experiment, and curated database, respectively. (c) Illustration of somatic mutations on PEG3 and APC genes in Chinese and TCGA data. (d) PEG3 and APC gene frequencies for Chinese and TCGA data (Chi-Squared test).

Seven significant mutated genes were identified, including 6 classical CRC genes (*TP53, APC, KRAS, FBXW7, PIK3CA* and *SMAD4* [5, 6]) and a novel CRC gene (PEG3). *PEG3* accounts for 9 (10.6%) of the patients compared with 2.7% mutation prevalence in TCGA data. Both of Illumina and Complete Genomics platforms are able to identify the somatic mutations of *PEG3* gene with similar mutation frequencies (3/29 and 6/56) for regularly mutated samples. Illustrations of the mutation sites are also shown in Supplementary Figure S1. Its aberrations on multiple levels, including gene expression, methylation and mutation have been reported in different types of tumors, including myeloma [22], ovarian cancers [23] and cholangiocarcinoma [24]. Moreover, expression of *PEG3* in colorectal patients shows statistically significant lower gene expression than matched controls from two independent studies using RNA-Seq and microarray platforms [20, 21]. Its reduced expression is also reported to be statistically significant associated with patients’ survival in various types of tumors [25]. *PEG3* acts as a tumor suppressor gene by binding and promoting the degradation of β-catenin, which will consequently inhibit Wnt Signaling [26]. Its molecular role is to interact with *SIAH1* and induce *TP53* mediated apoptosis or bind with *TRAF2* to initiate *NFKB* and *MAPK* pathways [27, 28].

A few other genes with mutation frequency greater than 5% were reported in this study and their association with CRC is revealed previously, such as *HMCN1* [29], *SYNE1 [10], NEB* [30], *OBSCN* [31], *MUC16, RYR2* [32] and *FAT4* [10]. Several genes showed medium mutation prevalence, including *ABCA13, ABCC10, CEP192, DCHS2, DNAH5, DYNC1H1, PTPRT, MYO16, LRRK2, DYNC2H1, FBN1, FCGBP, FLG, FREM2, PAPPA2, RP1L1, TTN, ZFHX4* and *ZNF717.* Their possible roles in CRC may be by interacting with canonical CRC genes in a network fashion as illustrated in Figure 3b.

Two genes display significantly different mutation frequencies between Chinese and TCGA data. The mutation rate of *APC* gene is 0.435 and 0.786 in Chinese and TCGA data (Figure 3c). This result is validated by previous Chinese exome-sequencing results (0.56 mutation frequency, p=0.0001 from Fisher’ exact test) [10]. *APC* gene carries a hotspot mutation Q1429 in Chinese population, differs from the hotspot mutation (R1450) in TCGA data (Figure 3d). Two novel hotspot mutations, namely R273C/H, R282W on *TP53* gene and R265C/H on *SMAD4* gene were shown in this dataset (Supplementary Figure S2). Additionally, *KRAS* and *PIK3CA* genes are less frequently mutated but not statistically significant in Chinese patients were also observed in this dataset.

Lastly, we used 1000 Genome Project and an in-house Chinese control dataset (1500 whole exome sequencing) to evaluate the possible systematic effects. It is shown that only 5.4% of somatic SNVs from our data were annotated in 1000 Genome Project, which is highly concordant with the TCGA results (6.8%). Only 2% of the somatic SNVs are found in the in-house data set, and most of them achieve very low mutation frequencies (Supplementary Table S7). This indicates the possible pathogenic nature of these variants.

### 2.4 Pathway analysis of mutated genes

We also investigate the mutation pattern on pathway level by mapping the somatic mutated genes onto the predefined cancer pathways [33, 34]. The method reported previously is used to characterize the pathway mutation frequencies [35]. Significantly mutated pathways (SMP) in Chinese population agree with TCGA results, including canonical pathways, such as Wnt/beta-catenin signaling, Cell Cycle/Apoptosis, MAPK signaling, TGF-beta signaling and PI(3)K signaling [5] as shown in Figure 4a [5]. It should be noted that *APC* pathway is identified as significantly mutated pathway in TCGA data, but not in our data. This is consistent with gene mutation frequency alteration results. Different pathway mutation frequency is observed, such as lower mutations rate on Wnt/beta-catenin signaling (44.7% versus 67.2%, p-value= 1.514E-5) and MAPK signaling (41.2% versus 51.2%, p-value=0.04485), and higher mutations rate on DNA damage control (64.7% versus 55.2%), Genome Integrity (62.4% versus 56.8%) and Cell Cycle/Apoptosis (65.9% versus 56.8%). Illustration of the pathway mutation frequency and associated genes are shown in Figure 4b and a report of the significantly mutated pathways is shown in Supplementary Table S8. These two platforms achieve quite high consistency in terms of the significant mutated pathways identification.

**Fig. 4.**
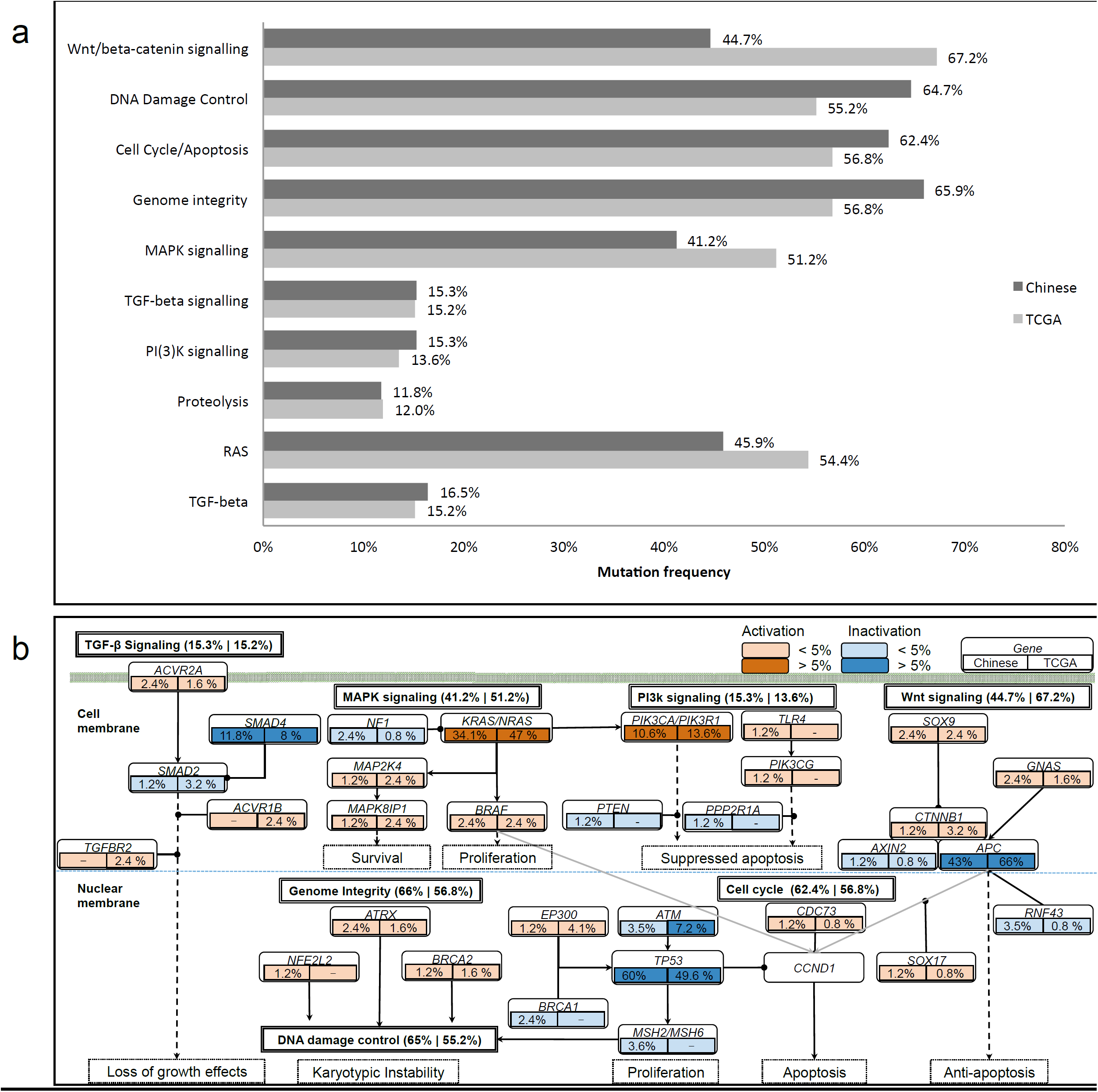
Illustration of mutated genes in pathways. (a). Comparison of significantly mutated pathways between Chinese and TCGA data (q-value<0.01). (b). Illustration of the mutated genes on pathway level. For each gene, their mutation frequency in Chinese and TCGA data are presented on the left and right box below each gene. The activation and inactivation frequencies >5% and <5% are denoted with respective colors as shown on the top right of the figure. The overall mutation frequencies for each pathway are described on the right of the pathway, coming with that of the Chinese and TCGA cohorts respectively.

## 3 DISCUSSION

We investigated the Chinese CRC tumorigenesis process by a whole exome sequencing in this study. We validated well-known somatic mutations in CRC, such as *TP53, APC, KRAS* and discovered a few high prevalent novel somatic mutations, including *PEG3* and *PTPRT.* Their mutation frequencies were also compared with that in TCGA data, and significant alternative frequency in *APC* or *PEG3* was observed. Pathway analysis of somatic mutations indicates a higher mutation frequency in cell cycle and lower mutation frequency in Wnt or MAPK signaling pathway.

The clinical characteristics association analyses were taken for genes with >5% mutation prevalence in regularly mutated samples, and the association results of *TP53, APC, KRAS, PEG3* or *PIK3CA* genes are shown in Table 1. *RP1L1* and *SYNE1* genes were significantly enriched in male and female patients. *PEG3* and *RYR2* displayed a significant higher association with early colorectal cancer (onset age≤45). Of note, mutations on *TP53* had a nearly two-fold preference in rectum (70.5%) and left colon (62.5%) than left colon (35.3%) (p-value=0.044). *APC* mutation frequency decreased with the increment of TNM stages, I+II (60.5%), III (26.3%) and IV (25%) (p-value = 0.004), and this pattern still persists when combining TNM III and TNM IV for this analysis. In contrast, TCGA data showed consistent mutation frequencies among TNM I+II (80%), III (76%) and IV (78.1%) patients (Supplementary Table S9). Our findings on alterations of the mutation frequency of *APC* in Chinese patients are consistent with the previous published results[36, 37]. Our identification on mutational pathways shows rates of Wnt/beta-catenin signaling in Chinese patients decrease from 0.651 (TNM I+II) to 0.262 (TNM III+IV), while mutational analysis of pathways of DNA damage control, genome integrity and cell Cycle/Apoptosis did not show a significant change. Meanwhile, our analysis of the mutation rate of TCGA data showed a consistence between TNM I+II (0.829) and TNM III+IV (0.793) stages for Wnt/beta-catenin signaling.

**Table 1.**
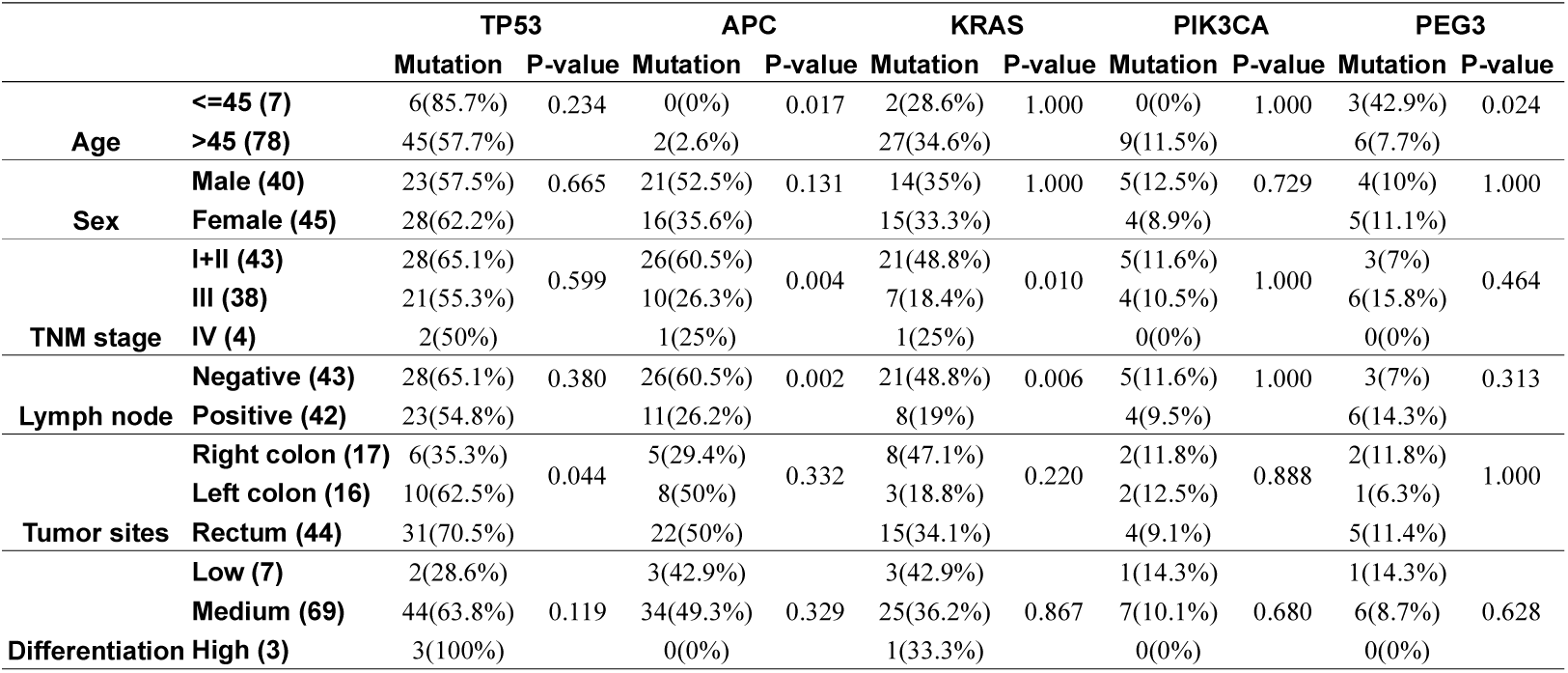
Clinical association with somatic mutated genes for non-hypermuated patients.

|  |  | TP53 |  | APC |  | KRAS |  | PIK3CA |  | PEG3 |  |
| --- | --- | --- | --- | --- | --- | --- | --- | --- | --- | --- | --- |
|  |  | Mutation | P-value | Mutation | P-value | Mutation | P-value | Mutation | P-value | Mutation | P-value |
| **Age** | **<=45 (7)** | 6(85.7%) | 0.234 | 0(0%) | 0.017 | 2(28.6%) | 1.000 | 0(0%) | 1.000 | 3(42.9%) | 0.024 |
|  | **>45 (78)** | 45(57.7%) |  | 2(2.6%) |  | 27(34.6%) |  | 9(11.5%) |  | 6(7.7%) |  |
| **Sex** | **Male (40)** | 23(57.5%) | 0.665 | 21(52.5%) | 0.131 | 14(35%) | 1.000 | 5(12.5%) | 0.729 | 4(10%) | 1.000 |
|  | **Female (45)** | 28(62.2%) |  | 16(35.6%) |  | 15(33.3%) |  | 4(8.9%) |  | 5(11.1%) |  |
| **TNM stage** | **I+II (43)** | 28(65.1%) | 0.599 | 26(60.5%) | 0.004 | 21(48.8%) | 0.010 | 5(11.6%) | 1.000 | 3(7%) | 0.464 |
|  | **III (38)** | 21(55.3%) |  | 10(26.3%) |  | 7(18.4%) |  | 4(10.5%) |  | 6(15.8%) |  |
|  | **IV (4)** | 2(50%) |  | 1(25%) |  | 1(25%) |  | 0(0%) |  | 0(0%) |  |
| **Lymph node** | **Negative (43)** | 28(65.1%) | 0.380 | 26(60.5%) | 0.002 | 21(48.8%) | 0.006 | 5(11.6%) | 1.000 | 3(7%) | 0.313 |
|  | **Positive (42)** | 23(54.8%) |  | 11(26.2%) |  | 8(19%) |  | 4(9.5%) |  | 6(14.3%) |  |
| **Tumor sites** | **Right colon (17)** | 6(35.3%) | 0.044 | 5(29.4%) | 0.332 | 8(47.1%) | 0.220 | 2(11.8%) | 0.888 | 2(11.8%) | 1.000 |
|  | **Left colon (16)** | 10(62.5%) |  | 8(50%) |  | 3(18.8%) |  | 2(12.5%) |  | 1(6.3%) |  |
|  | **Rectum (44)** | 31(70.5%) |  | 22(50%) |  | 15(34.1%) |  | 4(9.1%) |  | 5(11.4%) |  |
| **Differentiation** | **Low (7)** | 2(28.6%) |  | 3(42.9%) |  | 3(42.9%) |  | 1(14.3%) |  | 1(14.3%) |  |
|  | **Medium (69)** | 44(63.8%) | 0.119 | 34(49.3%) | 0.329 | 25(36.2%) | 0.867 | 7(10.1%) | 0.680 | 6(8.7%) | 0.628 |
|  | **High (3)** | 3(100%) |  | 0(0%) |  | 1(33.3%) |  | 0(0%) |  | 0(0%) |  |

It is widely accepted that sequential mutations on *APC*->*KRAS*->*TP53* is a key CRC development pathway [38–40]. The reduction of *APC* mutations rate in TNM III+IV patients undermine its importance in Chinese patients tumor progression. This will greatly shape our understanding about tumor progression, and may have direct clinical impact on Chinese CRC precision medicine management, for example, *APC* has been recommended as prognostic gene for survival prediction in Caucasian patients and our finding of lower mutation rate of *APC* may diminish its utility in Chinese patients [41]. The possible reason for lower *APC* mutation frequency in TNM III+IV tumor could be due to complementary mechanisms like *PEG3*, which has been reported to inhibit Wnt signaling by degrading β-catenin [26]. Another possible reason of decreased mutation frequency of *APC* and Wnt signaling in lymph node positive cancer may due to tumor cell eradication by immune cell, if these mutated genes are not essential for CRC maintenance.

In conclusion, we carried out the first large-scale (∼100 samples) whole exome-sequencing project in Chinese patients. *PEG3* has been identified as novel somatic mutated gene, and alternative mutation frequencies from TCGA data have been revealed (*APC* gene and WNT signaling). These newly identified somatic mutations on the canonical CRC genes worth further analysis. Taken together, the above study will be the genomics reference for the further colorectal cancer research in Chinese patients, and the raw sequences deposited at BIGD database (ID: CRA000626) will make that possible. The further updates on gene structural annotation, especially identification of novel exons, may change the results and derive interesting finding of the variants’ pathogenic feature [42, 43]. We are interested to see the results derived from the joint analysis of this study with other genomics data.

## 4 MATERIALS AND METHODS

### 4.1 Sample collection and preparation

During this study, 40 and 58 unrelated Chinese patients with colorectal cancer were recruited from Second Affiliated Hospital of Nanchang University and Wenzhou Medical University. They were referred for surgery from 2008 to 2010. These unrelated patients do not have a family history of colorectal cancer. All patients provided a written informed consent according to the research protocol approved by the Ethical Review Board at two hospitals. Fresh tissue samples were dissected and stored at liquid nitrogen. A description of the clinical characteristics is shown in Supplementary Table S1.

### 4.2 Illumina Exome Capture and Sequencing

Human genomic DNA was extracted from frozen tumor or matched colon tissue with QIAampDNA Mini kits (Qiagen). Covaris system is used to obtain the fragmented genomic DNA between 150 to 200bp. Adaptors were ligated to both ends of the fragments and adaptor-ligated templates were then purified using AgencourtAMPure SPRI beads. Whole-exome enrichment was performed by the SureSelect Human All Exon 51M kit (Agilent). Captured DNA libraries are sequenced by HiSeq 2000 Genome Analyzer (Illumina), resulting of 90 base paired-end reads.

### 4.3 Complete Genomics Exome Capture and Sequencing

Whole-exome sequencing was performed by a sequencing-by-ligation method [44]. Briefly, fragmented genomic DNA was generated by Covaris system with length between 200-400bp. After ligation and PCR amplification, the whole exome is enriched by Exome 59M kit (BGI). Then, a second round PCR amplification is performed. The resulted DNA fragments are prepared for Complete Genomics Black Bird Sequencer, and 30–35 base paired-end reads are finally obtained.

### 4.4 Bioinformatics analysis

#### 4.4.1 Illumina variant detection

The following pipeline was carried out with default parameters unless explicitly described. First, clean reads were obtained by removing adapter reads and low quality reads (≥10% of the bases are N, or ≥50% of the bases with Phred score ≤5). Burrows-Wheeler Aligner (BWA) was used to align the clean reads to the human reference genome (UCSC Genome Browser hg19) with parameters “*-o 1 -e 50 –m 100000 -t 4 -i 15 -q 10 -I”* [45]. SAMtools was used to convert the SAM-formatted alignment results to BAM-formatted alignment files. Local read alignment re-calibration was performed by Genome Analysis Toolkit (GATK IndelRealigner) [46]. Finally, Picard toolkit was used to mark duplicates. [47]. **Somatic SNV detection**: Mutect v1.1.4 was used to compare tumor BAM files against their matched control BAM files for somatic single nucleotide variants (SNVs) identification [48]. They are filtered with the following requirements: *minimum coverage* ≥ *10X, mutation allele fraction* ≥*10% and* ≥*5 reads*. **Somatic Indel detection**: GATK was used to detect somatic InDels with following parameters: *minCoverage=6, minNormalCoverage=4, minFraction=0.3*. False positive InDels were finally removed by in-house scripts.

#### 4.4.2 Complete Genomics variant detection

The resulting mate-paired reads were aligned to the reference genome (hg19) and variants are called by the reported methods [49].

### 4.5 Statistical analysis

Qualitative variables were compared by Fisher’s exact test. *T-test* was used for normal distributed data comparison, and Wilcoxon rank test was used for non-normal distributed data. All of the statistical analyses were performed in R or Bioconductor environment.

## ABBREVIATIONS

CRC: colorectal cancer
TCGA: The Cancer Genome Atlas
SNV: Single Nucleotide variation
InDel: Insertion and deletion

## AUTHOR CONTRIBUTIONS

JD and XDF designed the project, MY initiated the project and collected the samples, CY, ZL, YZ, XCL and WL performed the bioinformatics analysis; ZL, YC, YZ, XDF and JD wrote the paper. All of the authors agree the manuscript.

## ACKNOWLEDGEMENTS

Thanks are given to Rongfa Yuan, Jianghua Shao and Chunmei Piao who have helped us to collect the patient samples, and Prasanth Bhatt who proofread this manuscript.

## REFERENCES

1. Chen W, Zheng R, Baade PD, Zhang S, Zeng H, Bray F, Jemal A, Yu XQ, He J. Cancer statistics in China, 2015. CA Cancer J Clin. 2016; 66: 115–32. doi: 10.3322/caac.21338.

2. Siegel RL, Miller KD, Fedewa SA, Ahnen DJ, Meester RGS, Barzi A, Jemal A. Colorectal cancer statistics, 2017. CA: A Cancer Journal for Clinicians. 2017; 67: 177–93. doi: 10.3322/caac.21395.

3. Xu AG, Yu ZJ, Jiang B, Wang XY, Zhong XH, Liu JH, Lou QY, Gan AH. Colorectal cancer in Guangdong Province of China: a demographic and anatomic survey. World J Gastroenterol. 2010; 16: 960–5. doi: 10.3748/wjg.v16.i8.960

4. Wood LD, Parsons DW, Jones S, Lin J, Sjöblom T, Leary RJ, Shen D, Boca SM, Barber T, Ptak J, Silliman N, Szabo S, Dezso Z, et al. The Genomic Landscapes of Human Breast and Colorectal Cancers. Science. 2007; 318: 1108–13. doi: 10.1126/science.1145720.

5. Cancer Genome Atlas N. Comprehensive molecular characterization of human colon and rectal cancer. Nature. 2012; 487: 330–7. doi: 10.1038/nature11252.

6. Seshagiri S, Stawiski EW, Durinck S, Modrusan Z, Storm EE, Conboy CB, Chaudhuri S, Guan Y, Janakiraman V, Jaiswal BS. Recurrent R-spondin fusions in colon cancer. Nature. 2012; 488: 660–4. doi: 10.1038/nature11282

7. Guda K, Veigl ML, Varadan V, Nosrati A, Ravi L, Lutterbaugh J, Beard L, Willson JKV, Sedwick WD, Wang ZJ, Molyneaux N, Miron A, Adams MD, et al. Novel recurrently mutated genes in African American colon cancers. Proceedings of the National Academy of Sciences of the United States of America. 2015; 112: 1149–54. doi: 10.1073/pnas.1417064112.

8. Nagahashi M, Wakai T, Shimada Y, Ichikawa H, Kameyama H, Kobayashi T, Sakata J, Yagi R, Sato N, Kitagawa Y, Uetake H, Yoshida K, Oki E, et al. Genomic landscape of colorectal cancer in Japan: clinical implications of comprehensive genomic sequencing for precision medicine. Genome Medicine. 2016; 8: 136. doi: 10.1186/s13073-016-0387-8.

9. Ashktorab H, Mokarram P, Azimi H, Olumi H, Varma S, Nickerson ML, Brim H. Targeted exome sequencing reveals distinct pathogenic variants in Iranians with colorectal cancer. Oncotarget. 2017; 8: 7852–66. doi: 10.18632/oncotarget.13977.

10. Yu J, Wu WK, Li X, He J, Li XX, Ng SS, Yu C, Gao Z, Yang J, Li M, Wang Q, Liang Q, Pan Y, et al. Novel recurrently mutated genes and a prognostic mutation signature in colorectal cancer. Gut. 2015; 64: 636–45. doi: 10.1136/gutjnl-2013-306620.

11. Giannakis M, Mu XJ, Shukla SA, Qian ZR, Cohen O, Nishihara R, Bahl S, Cao Y, Amin-Mansour A, Yamauchi M, Sukawa Y, Stewart C, Rosenberg M, et al. Genomic Correlates of Immune-Cell Infiltrates in Colorectal Carcinoma (vol 15, pg 857, 2016). Cell Reports. 2016; 17: 1206-. doi: 10.1016/j.celrep.2016.10.009.

12. Lam HYK, Clark MJ, Chen R, Chen R, Natsoulis G, O’Huallachain M, Dewey FE, Habegger L, Ashley EA, Gerstein MB, Butte AJ, Ji HP, Snyder M. Performance comparison of whole-genome sequencing platforms. Nature Biotechnology. 2012; 30: 78–U118. doi: 10.1038/nbt.2065.

13. Seshagiri S, Stawiski EW, Durinck S, Modrusan Z, Storm EE, Conboy CB, Chaudhuri S, Guan YH, Janakiraman V, Jaiswal BS, Guillory J, Ha C, Dijkgraaf GJP, et al. Recurrent R-spondin fusions in colon cancer. Nature. 2012; 488: 660-+. doi: 10.1038/nature11282.

14. Giannakis M, Mu XJ, Shukla SA, Qian ZR, Cohen O, Nishihara R, Bahl S, Cao Y, Amin-Mansour A, Yamauchi M, Sukawa Y, Stewart C, Rosenberg M, et al. Genomic Correlates of Immune-Cell Infiltrates in Colorectal Carcinoma. Cell Rep. 2016; 17: 1206. doi: 10.1016/j.celrep.2016.10.009.

15. Yu C, Yu J, Yao X, Wu WK, Lu Y, Tang S, Li X, Bao L, Li X, Hou Y, Wu R, Jian M, Chen R, et al. Discovery of biclonal origin and a novel oncogene SLC12A5 in colon cancer by single-cell sequencing. Cell Res. 2014; 24: 701–12. doi: 10.1038/cr.2014.43.

16. Wu H, Zhang XY, Hu Z, Hou Q, Zhang H, Li Y, Li S, Yue J, Jiang Z, Weissman SM, Pan X, Ju BG, Wu S. Evolution and heterogeneity of non-hereditary colorectal cancer revealed by single-cell exome sequencing. Oncogene. 2016. doi: 10.1038/onc.2016.438.

17. Tokheim CJ, Papadopoulos N, Kinzler KW, Vogelstein B, Karchin R. Evaluating the evaluation of cancer driver genes. Proc Natl Acad Sci U S A. 2016; 113: 14330–5. doi: 10.1073/pnas.1616440113.

18. Hofree M, Carter H, Kreisberg JF, Bandyopadhyay S, Mischel PS, Friend S, Ideker T. Challenges in identifying cancer genes by analysis of exome sequencing data. Nat Commun. 2016; 7: 12096. doi: 10.1038/ncomms12096.

19. Lawrence MS, Stojanov P, Polak P, Kryukov GV, Cibulskis K, Sivachenko A, Carter SL, Stewart C, Mermel CH, Roberts SA, Kiezun A, Hammerman PS, McKenna A, et al. Mutational heterogeneity in cancer and the search for new cancer-associated genes. Nature. 2013; 499: 214–8. doi: 10.1038/nature12213.

20. Kim SK, Kim SY, Kim JH, Roh SA, Cho DH, Kim YS, Kim JC. A nineteen gene-based risk score classifier predicts prognosis of colorectal cancer patients. Mol Oncol. 2014; 8: 1653–66. doi: 10.1016/j.molonc.2014.06.016.

21. Matsuyama T, Ishikawa T, Mogushi K, Yoshida T, Iida S, Uetake H, Mizushima H, Tanaka H, Sugihara K. MUC12 mRNA expression is an independent marker of prognosis in stage II and stage III colorectal cancer. Int J Cancer. 2010; 127: 2292–9. doi: 10.1002/ijc.25256.

22. Bolli N, Avet-Loiseau H, Wedge DC, Van Loo P, Alexandrov LB, Martincorena I, Dawson KJ, Iorio F, Nik-Zainal S, Bignell GR, Hinton JW, Li Y, Tubio JM, et al. Heterogeneity of genomic evolution and mutational profiles in multiple myeloma. Nat Commun. 2014; 5: 2997. doi: 10.1038/ncomms3997.

23. Feng W, Marquez RT, Lu Z, Liu J, Lu KH, Issa J-PJ, Fishman DM, Yu Y, Bast RC. Imprinted tumor suppressor genes ARHI and PEG3 are the most frequently down-regulated in human ovarian cancers by loss of heterozygosity and promoter methylation. Cancer. 2008; 112: 1489–502. doi: 10.1002/cncr.23323.

24. Ong CK, Subimerb C, Pairojkul C, Wongkham S, Cutcutache I, Yu W, McPherson JR, Allen GE, Ng CC, Wong BH, Myint SS, Rajasegaran V, Heng HL, et al. Exome sequencing of liver fluke-associated cholangiocarcinoma. Nat Genet. 2012; 44: 690–3. doi: 10.1038/ng.2273.

25. Li M, Sun Q, Wang X. Transcriptional landscape of human cancers. Oncotarget. 2017; 8: 34534–51. doi: 10.18632/oncotarget.15837.

26. Jiang X, Yu Y, Yang HW, Agar NY, Frado L, Johnson MD. The imprinted gene PEG3 inhibits Wnt signaling and regulates glioma growth. J Biol Chem. 2010; 285: 8472–80. doi: 10.1074/jbc.M109.069450.

27. Relaix F, Wei XJ, Li W, Pan JJ, Lin YH, Bowtell DD, Sassoon DA, Wu XW. Pw1/Peg3 is a potential cell death mediator and cooperates with Siah1a in p53-mediated apoptosis. Proceedings of the National Academy of Sciences of the United States of America. 2000; 97: 2105–10. doi: 10.1073/Pnas.040378897.

28. Relaix F, Wei XJ, Wu X, Sassoon DA. Peg3/Pw1 is an imprinted gene involved in the TNF-NFkappaB signal transduction pathway. Nat Genet. 1998; 18: 287–91. doi: 10.1038/ng0398-287.

29. Lee SH, Je EM, Yoo NJ, Lee SH. HMCN1, a cell polarity-related gene, is somatically mutated in gastric and colorectal cancers. Pathol Oncol Res. 2015; 21: 847–8. doi: 10.1007/s12253-014-9809-3.

30. Kortum KM, Langer C, Monge J, Bruins L, Zhu YX, Shi CX, Jedlowski P, Egan JB, Ojha J, Bullinger L, Kull M, Ahmann G, Rasche L, et al. Longitudinal analysis of 25 sequential sample-pairs using a custom multiple myeloma mutation sequencing panel (M(3)P). Ann Hematol. 2015; 94: 1205–11. doi: 10.1007/s00277-015-2344-9.

31. Balakrishnan A, Bleeker FE, Lamba S, Rodolfo M, Daniotti M, Scarpa A, van Tilborg AA, Leenstra S, Zanon C, Bardelli A. Novel somatic and germline mutations in cancer candidate genes in glioblastoma, melanoma, and pancreatic carcinoma. Cancer Res. 2007; 67: 3545–50. doi: 10.1158/0008-5472.CAN-07-0065.

32. Bueno R, Stawiski EW, Goldstein LD, Durinck S, De Rienzo A, Modrusan Z, Gnad F, Nguyen TT, Jaiswal BS, Chirieac LR, Sciaranghella D, Dao N, Gustafson CE, et al. Comprehensive genomic analysis of malignant pleural mesothelioma identifies recurrent mutations, gene fusions and splicing alterations. Nat Genet. 2016; 48: 407–16. doi: 10.1038/ng.3520.

33. Vogelstein B, Papadopoulos N, Velculescu VE, Zhou S, Diaz LA, Jr., Kinzler KW. Cancer genome landscapes. Science. 2013; 339: 1546–58. doi: 10.1126/science.1235122.

34. Kandoth C, McLellan MD, Vandin F, Ye K, Niu B, Lu C, Xie M, Zhang Q, McMichael JF, Wyczalkowski MA, Leiserson MDM, Miller CA, Welch JS, et al. Mutational landscape and significance across 12 major cancer types. Nature. 2013; 502: 333–9. doi: 10.1038/nature12634.

35. Li X, Wu WK, Xing R, Wong SH, Liu Y, Fang X, Zhang Y, Wang M, Wang J, Li L, Zhou Y, Tang S, Peng S, et al. Distinct Subtypes of Gastric Cancer Defined by Molecular Characterization Include Novel Mutational Signatures with Prognostic Capability. Cancer Res. 2016; 76: 1724–32. doi: 10.1158/0008-5472.CAN-15-2443.

36. Liu X, Shan X, Friedl W, Uhlhaas S, Propping P, Li J, Wang Y. May the APC gene somatic mutations in tumor tissues influence the clinical features of Chinese sporadic colorectal cancers? Acta Oncologica. 2007; 46: 757–62. doi: 10.1080/02841860600996439.

37. Ko JM, Cheung MH, Kwan MW, Wong CM, Lau KW, Tang CM, Lung ML. Genomic instability and alterations in Apc, Mcc and Dcc in Hong Kong patients with colorectal carcinoma. Int J Cancer. 1999; 84: 404–9. doi: 10.1002/(SICI)1097-0215(19990820)84:4<404::AID-IJC13>3.0.CO;2-L

38. Markowitz SD, Bertagnolli MM. Molecular origins of cancer: Molecular basis of colorectal cancer. N Engl J Med. 2009; 361: 2449–60. doi: 10.1056/NEJMra0804588.

39. Jones S, Chen WD, Parmigiani G, Diehl F, Beerenwinkel N, Antal T, Traulsen A, Nowak MA, Siegel C, Velculescu VE, Kinzler KW, Vogelstein B, Willis J, et al. Comparative lesion sequencing provides insights into tumor evolution. Proc Natl Acad Sci U S A. 2008; 105: 4283–8. doi: 10.1073/pnas.0712345105.

40. Fearon ER, Vogelstein B. A genetic model for colorectal tumorigenesis. Cell. 1990; 61: 759–67. doi: 10.1016/0092-8674(90)90186-I

41. Schell MJ, Yang M, Teer JK, Lo FY, Madan A, Coppola D, Monteiro ANA, Nebozhyn MV, Yue B, Loboda A, Bien-Willner GA, Greenawalt DM, Yeatman TJ. A multigene mutation classification of 468 colorectal cancers reveals a prognostic role for APC. Nature Communications. 2016; 7: 11743. doi: 10.1038/ncomms11743

42. Steward CA, Parker APJ, Minassian BA, Sisodiya SM, Frankish A, Harrow J. Genome annotation for clinical genomic diagnostics: strengths and weaknesses. Genome Med. 2017; 9: 49. doi: 10.1186/s13073-017-0441-1.

43. Liu Z, Ma H, Goryanin I. A semi-automated genome annotation comparison and integration scheme. BMC Bioinformatics. 2013; 14: 172. doi: 10.1186/1471-2105-14-172.

44. Drmanac R, Sparks AB, Callow MJ, Halpern AL, Burns NL, Kermani BG, Carnevali P, Nazarenko I, Nilsen GB, Yeung G, Dahl F, Fernandez A, Staker B, et al. Human genome sequencing using unchained base reads on self-assembling DNA nanoarrays. Science. 2010; 327: 78–81. doi: 10.1126/science.1181498.

45. Li H, Durbin R. Fast and accurate short read alignment with Burrows-Wheeler transform. Bioinformatics. 2009; 25: 1754–60. doi: 10.1093/bioinformatics/btp324.

46. Li H, Handsaker B, Wysoker A, Fennell T, Ruan J, Homer N, Marth G, Abecasis G, Durbin R, Genome Project Data Processing S. The Sequence Alignment/Map format and SAMtools. Bioinformatics. 2009; 25: 2078–9. doi: 10.1093/bioinformatics/btp352.

47. McKenna A, Hanna M, Banks E, Sivachenko A, Cibulskis K, Kernytsky A, Garimella K, Altshuler D, Gabriel S, Daly M, DePristo MA. The Genome Analysis Toolkit: a MapReduce framework for analyzing next-generation DNA sequencing data. Genome Res. 2010; 20: 1297–303. doi: 10.1101/gr.107524.110.

48. Cibulskis K, Lawrence MS, Carter SL, Sivachenko A, Jaffe D, Sougnez C, Gabriel S, Meyerson M, Lander ES, Getz G. Sensitive detection of somatic point mutations in impure and heterogeneous cancer samples. Nat Biotechnol. 2013; 31: 213–9. doi: 10.1038/nbt.2514.

49. Carnevali P, Baccash J, Halpern AL, Nazarenko I, Nilsen GB, Pant KP, Ebert JC, Brownley A, Morenzoni M, Karpinchyk V, Martin B, Ballinger DG, Drmanac R. Computational techniques for human genome resequencing using mated gapped reads. J Comput Biol. 2012; 19: 279–92. doi: 10.1089/cmb.2011.0201.

