## Supplementary Materials for "The landscape of somatic mutation in sporadic Chinese colorectal cancer"

**Appendix A Clustering of mutation spectrums**

We performed log ratio transformation of the six mutation type data, and then used hierarchical clustering with Manhattan distance and centroid linkage for clustering. Silhouette plot is utilized to choose the best number of clusters on the basis of average silhouette width. The average silhouette plot of Chinese data is maximized when the number of clusters equals to 2 (Figure 1). The clustering results of TCGA data also indicate two clusters (Figure 2). The clustering results indicate that samples in Cluster 1 correspond perfectly to hypermutated samples. TCGA data also shows similar pattern and all of 33 the samples in Cluster 2 are hypermutated samples, only 4 hypermutated samples reside in Cluster 1. The results of Chinese and TCGA data are shown in Figure 3 and 4.

**Figure 1 Silhouette plot of the Chinese data**

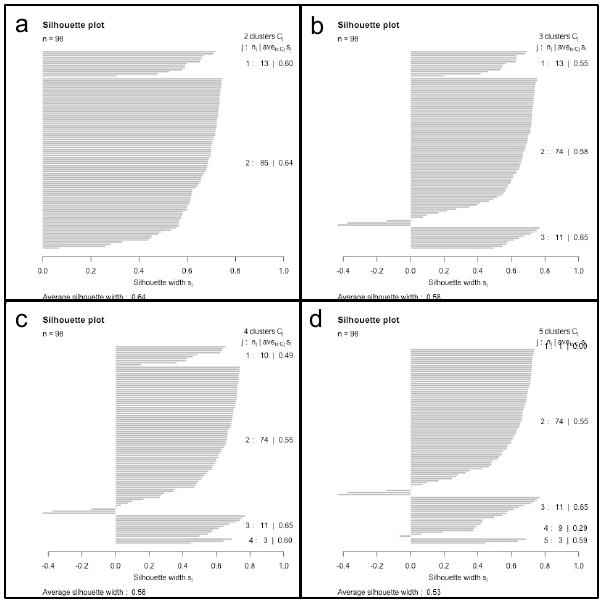

**(a), (b), (c) and (d) represent silhouette plot of 2, 3, 4 and 5 clusters of Chinese CRC data based on six mutation types**

**Figure 2 Silhouette plot of the TCGA data**

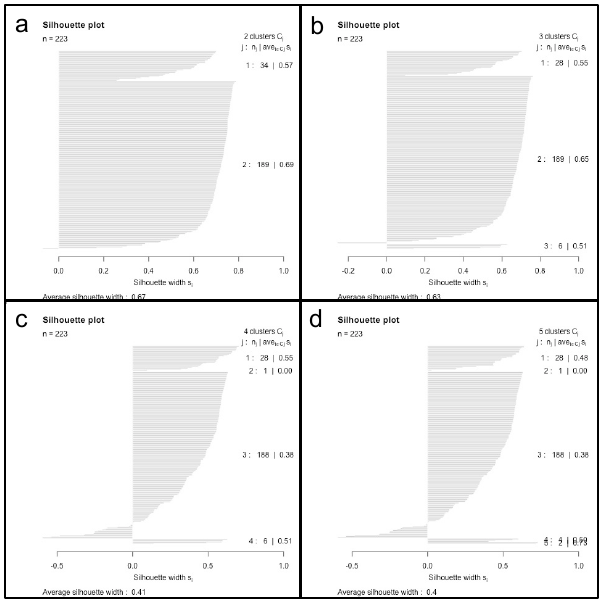

**(a), (b), (c) and (d) represent silhouette plot of 2, 3, 4 and 5 clusters of TCGA CRC data based on six mutation types**

**Figure 3 Clustering results of Chinese data**

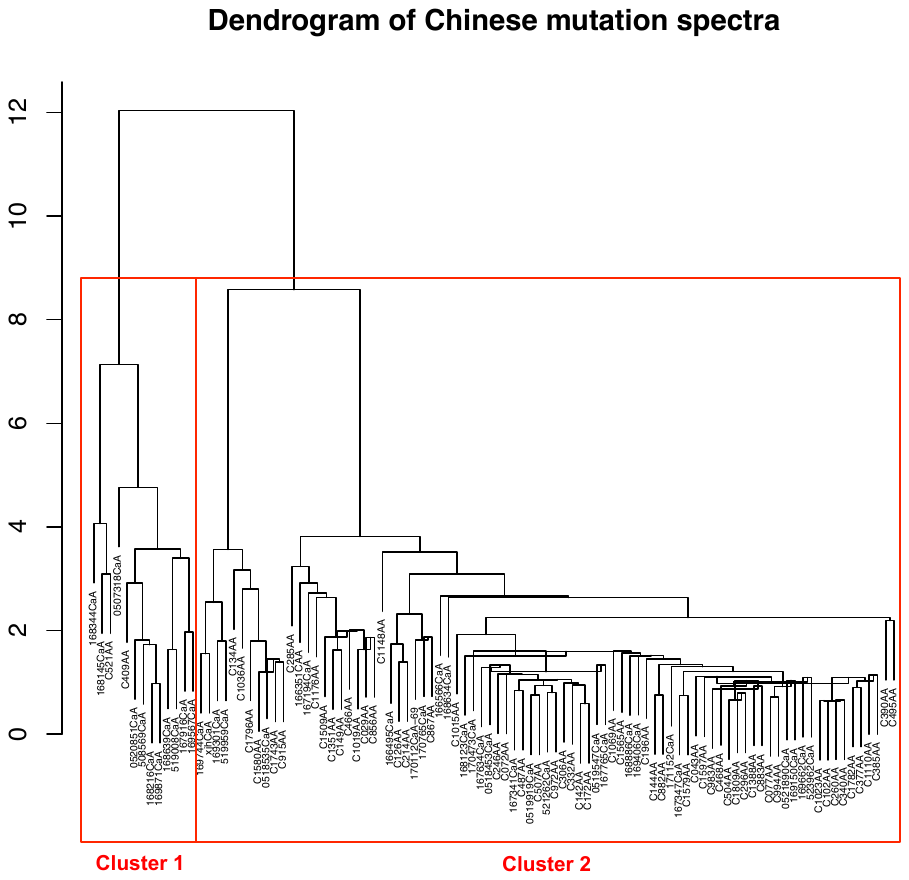

**Figure 4 Silhouette plot of the TCGA data**

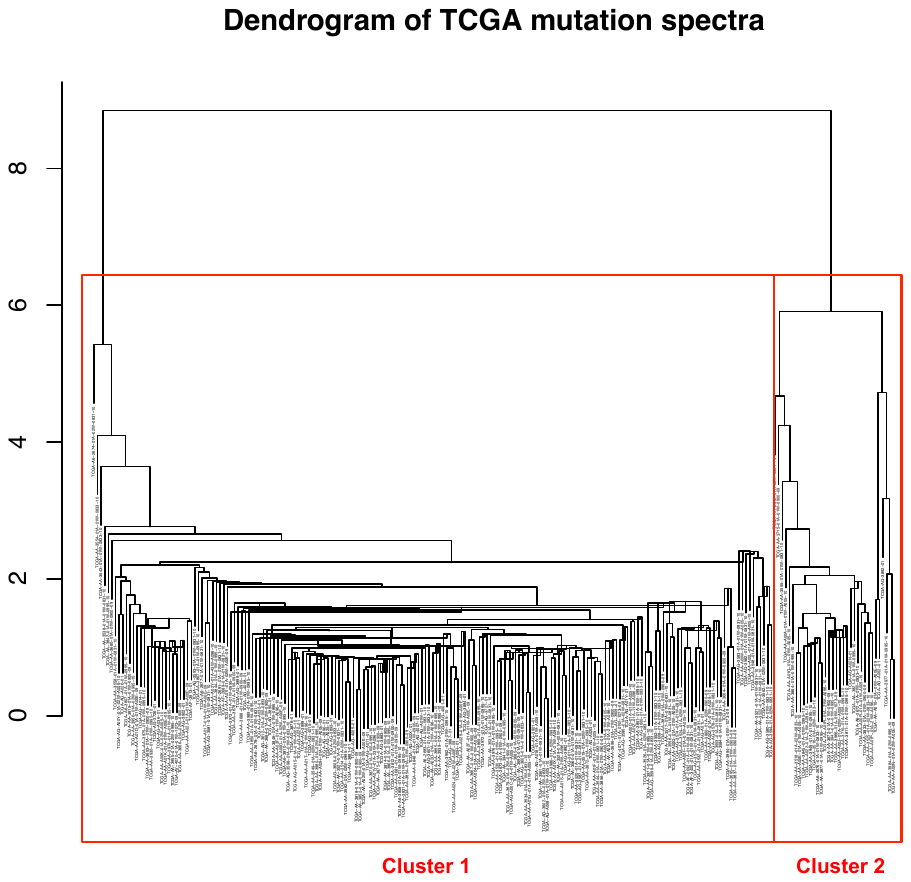
