## Supplementary Materials for "The landscape of somatic mutation in sporadic Chinese colorectal cancer"

**Supplementary Figure S1 Illustration of somatic mutations on PEG3**

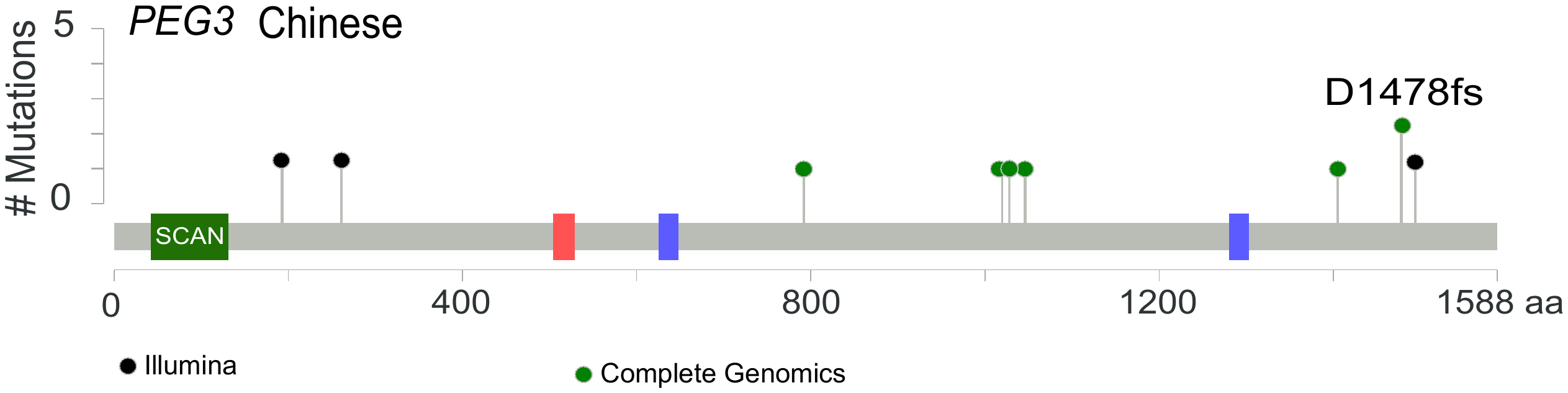

**Supplementary Figure S2 Illustration of somatic mutations on canonical CRC genes**

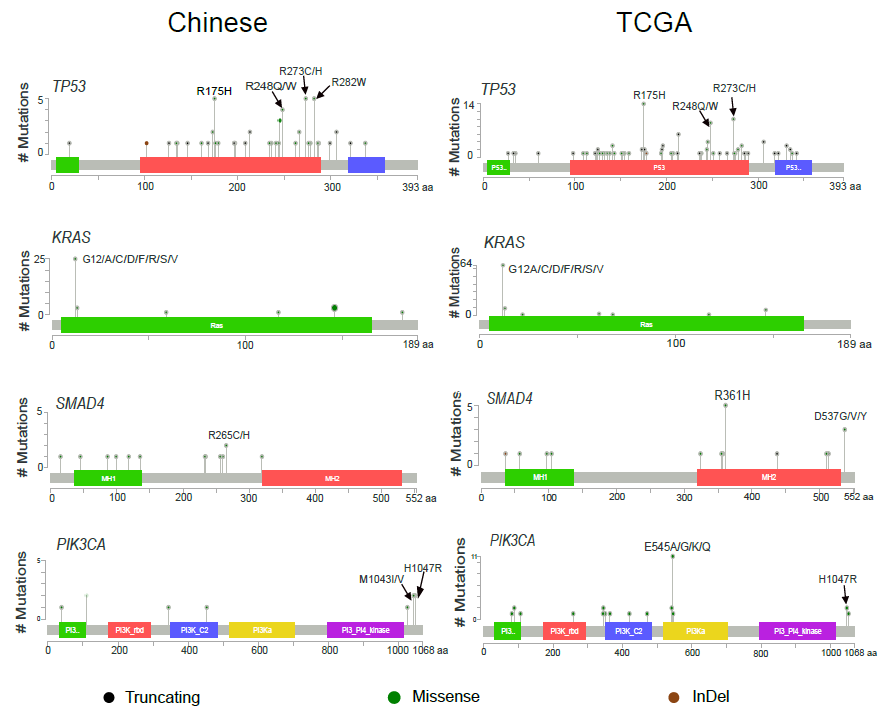
